# Role of SHIP2 in cell repulsion regulated by Eph receptor and ephrin signaling

**DOI:** 10.1101/872846

**Authors:** Tim G. Ashlin, Zhonglin Wu, Qiling Xu, David G. Wilkinson

## Abstract

Previous studies have found that activation of EphB2 and ephrinB1 that drives cell segregation leads to phosphorylation of the phosphoinositide phosphatase SHIP2 downstream of forward (EphB2) but not reverse (ephrinB1) signaling. We have analysed whether SHIP2 interacts with EphB2 and contributes to cell responses to EphB2-ephrinB1 signaling. We confirm that EphB2 activation leads to SHIP2 phosphorylation on Y1135 and find that they interact through the SH2 domain of SHIP2. There is thus a distinct mode of interaction from EphA2, which binds SHIP2 via its SAM domain. Knockdown of SHIP2 in EphB2 cells leads to decreased segregation from ephrinB1 cells, and a decrease in the repulsion response of EphB2 cells. SHIP2 knockdown in ephrinB1 cells also decreases their repulsion response, but does not disrupt segregation which is largely driven by forward signaling. These findings show that activation of EphB2 leads to recruitment and phosphorylation of SHIP2, and that SHIP2 contributes to cell repulsion responses that underlie cell segregation.

## INTRODUCTION

Eph receptor and ephrin signaling has a major role in the maintenance of tissue organisation in vertebrates by triggering cell responses that drive the segregation of distinct cell populations. The segregation of cells underlies the sharpening of borders and inhibition of intermingling between distinct tissues, or subdivisions within tissues (Batlle and Wilkinson, 2012; Dahmann et al., 2011; Fagotto, 2014). Eph receptor tyrosine kinases comprise a large family of receptors which mediate cell contact dependent signaling upon interacting with membrane-bound ephrins. These interactions are promiscuous but are largely subdivided into two classes: with some exceptions, EphAs bind to GPI-anchored ephrinA proteins, and EphBs to transmembrane ephrinB proteins (Gale et al., 1996). Upon binding of Eph to ephrin, both components become clustered and can transduce signals (Egea and Klein, 2007; Pasquale, 2008). The activation of Eph receptors leads to ‘forward’ signaling, in part through tyrosine kinase dependent pathways. Ephrins mediate ‘reverse’ signaling, which for ephrinB proteins involves phosphorylation of tyrosine residues by cytoplasmic tyrosine kinases (Egea and Klein, 2007; Pasquale, 2008). Since Eph receptors and ephrins that have high affinity are expressed in complementary regions (Gale et al., 1996; Rohani et al., 2014), bi-directional activation occurs at the interface.

Eph receptor and ephrin signaling can drive cell segregation by regulating the strength of cell adhesion, repulsion and/or cortical tension at the heterotypic interface where strong activation occurs (Batlle and Wilkinson, 2012; Cayuso et al., 2015; Fagotto et al., 2014). Eph receptor activation can decrease adhesion through local cleavage of cadherins (Solanas et al., 2011) and by modulation of cadherin clustering (Fagotto et al., 2013). This leads to decreased adhesion at heterotypic compared with homotypic contacts that can potentially drive cell segregation (Steinberg, 2007). In another mechanism, Eph receptor and ephrin activation triggers cell repulsion in which there is a collapse response and change in direction of movement to move away from each other (Astin et al., 2010; Poliakov et al., 2008; Rohani et al., 2011; Taylor et al., 2017). Such heterotypic repulsion can drive segregation of migratory cells (Taylor et al., 2017), and requires bi-directional repulsion that prevents each population from mingling into the other (Canty et al., 2017; Rohani et al., 2014; Wu et al., 2019). Forward signaling has a dominant role as it triggers strong repulsion (O’Neill et al., 2016; Rohani et al., 2011; Rohani et al., 2014; Taylor et al., 2017), but reverse signaling can also contribute (Wu et al., 2019). A further mechanism is important in epithelial tissues in which Eph receptor activation leads to actomyosin contraction that increases cortical tension at the interface (Calzolari et al., 2014; Cayuso et al., 2019; O’Neill et al., 2016; Voltes et al., 2019). This increase in cortical tension can drive cell segregation and act as a barrier to intermingling between Eph receptor and ephrin-expressing cells (Canty et al., 2017). Recent studies suggest that increased tension or cell repulsion at the heterotypic interface are more efficient mechanisms for cell segregation and border sharpening than differential adhesion (Canty et al., 2017; Taylor et al., 2017).

At present, there is a partial understanding of the mechanisms by which Eph receptor and ephrin signaling leads to cell repulsion. A prominent role is played by pathways that modulate the activity of Rho family GTPases that regulate polymerisation of the actin cytoskeleton and actomyosin contraction (Kania and Klein, 2016; Klein, 2012; Pasquale, 2008). These pathways are mainly activated by kinase-dependent tyrosine phosphorylation that recruits SH2 domain proteins (Kania and Klein, 2016; Klein, 2012; Pasquale, 2008), but can also involve proteins that constitutively interact, for example PDZ domain proteins that bind to the C-terminus of Eph receptors and ephrinBs (Cayuso et al., 2019; Tanaka et al., 2003). An important role is also played by mechanisms that regulate endocytosis. Interactions of Eph receptors and ephrins can potentially cause adhesion of cells as they have high affinity for each other and form multimeric complexes at the site of cell-cell contact. Insights into how the adhesive interaction does not interfere with repulsion came from studies showing that upon activation, EphB-ephrinB complexes are trans-endocytosed into the Eph- and/or ephrin-expressing cell (forward and reverse endocytosis, respectively) (Marston et al., 2003; Zimmer et al., 2003). By removing Eph-ephrin complexes from the cell surface, trans-endocytosis enables the cells to disengage concommitant with repulsion

Further potential mediators of cell responses to Eph receptor and ephrin signaling have been identified in HEK293 cell lines with stable overexpression of EphB2 or ephrinB1, for brevity termed EphB2 and ephrinB1 cells (Poliakov et al., 2008). When mixed in cell culture, EphB2 and ephrinB1 cells segregate from each other through heterotypic cell repulsion (Taylor et al., 2017). Targets of EphB2 and ephrinB1 activation were identified by use of SILAC to detect proteins with increased or decreased tyrosine phosphorylation (Jorgensen et al., 2009). This found many proteins with altered tyrosine phosphorylation downstream of EphB2 and/or ephrinB1 activation, which include regulators of cell polarity, cell adhesion and actin remodelling. One of the targets is the phosphoinositide phosphatase SHIP2 (INPPL1), which has increased phosphorylation of Y986 and Y1135 in response to forward but not reverse signaling. Furthermore, an siRNA screen of phosphorylation targets suggested that knockdown of SHIP2 decreases segregation of EphB2 and ephrinB1 cells (Jorgensen et al., 2009). Since phosphoinositides modulated by SHIP2 regulate endocytosis (Boucrot et al., 2015; Nakatsu et al., 2010; Posor et al., 2015; Zhuang et al., 2007), and the polarity and migration of cells (Charest and Firtel, 2006; Elong Edimo et al., 2016; Kato et al., 2012; Krause and Gautreau, 2014; Lam et al., 2012; Venkatareddy et al., 2011; Yu et al., 2010), these studies suggest that SHIP2 is involved in cell responses to EphB2 activation. SHIP2 has been shown to bind directly to EphA2 through interaction of their SAM domains, and to inhibit forward endocytosis of EphA2 (Zhuang et al., 2007). However, structural studies and binding assays find that SHIP2 cannot bind to EphB2 through its SAM domain (Lee et al., 2012; Wang et al., 2018), raising the question of whether there is a direct interaction between SHIP2 and EphB2.

We set out to address whether SHIP2 interacts with EphB2 and regulates cell responses to forward signaling. We confirm that SHIP2 is phosphorylated during forward signaling and find that it interacts with activated EphB2. Unlike the relationship with EphA2, SHIP2 interacts with EphB2 via its SH2 domain, and a change in receptor endocytosis is not detected after knockdown of SHIP2. Rather, SHIP2 knockdown leads to a decrease in cell repulsion responses to both forward and reverse signaling, and thus contributes to polarised cell migration that can drive cell segregation.

## RESULTS

### Activated EphB2 binds and increases phosphorylation of SHIP2

Phosphoproteomic analyses suggested that SHIP2 is phosphorylated on Y1135 during EphB2 (forward) but not ephrinB1 (reverse) signaling (Jorgensen et al., 2009). In order to verify this result, we activated forward and reverse signaling separately by incubating EphB2 cells with clustered ephrinB1-Fc or ephrinB1 cells with clustered EphB2-Fc, respectively. Phosphorylation of SHIP2 on Y1135 was found to occur upon EphB2 activation but not ephrinB1 activation (Fig.1A).

**Figure 1.**
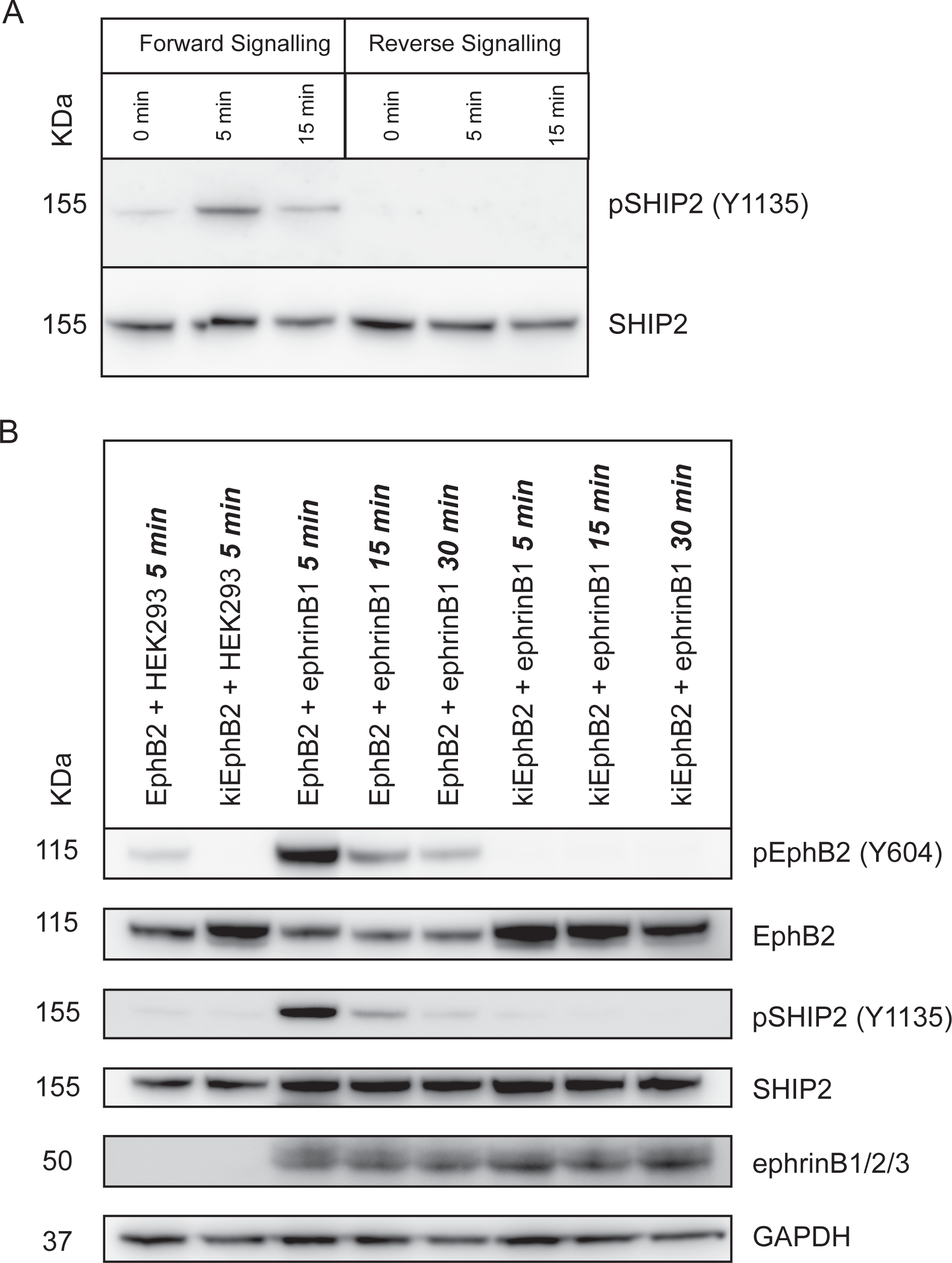
EphB2 activation leads to SHIP2 phosphorylation. Experiments were carried out to analyse phosphorylation of SHIP2 at Y1135 after activation of EphB2 and/or ephrinB1. (A): EphB2 cells were incubated with soluble ephrinB1-Fc or ephrinB1 cells were incubated with soluble EphB2-Fc for 0, 5 and 15 min. The cells were lysed and phosphorylation of SHIP2 at Y1135 was measured by western blotting. Phosphorylation occurs after forward but not reverse signaling. (B): To understand the temporal relationship between EphB2 activation and SHIP2 Y1135 phosphorylation, a cell mixing assay was used. EphB2 or kiEphB2 cells were mixed with ephrinB1 cells, briefly centrifuged to force the cells into contact and incubated for 5, 15 or 30 min. EphB2 activation (phosphorylation on Y604) and phosphorylation of SHIP2 on Y1135 have a similar time course, and were not observed when a kinase inactive mutant of EphB2 was used.

Since SHIP2 phosphorylation occurs only in EphB2 cells, we were able to determine the temporal relationship between SHIP2 phosphorylation and contact-dependent activation that occurs following mixing of EphB2 and ephrinB1 cells. We used an assay in which EphB2 is synchronously activated by ephrinB1: EphB2 cells and ephrinB1 cells are mixed in a tube, briefly centrifuged to force cells into contact, and then cells lysed for Western blot analysis at different time intervals (Wu et al., 2019). To reveal EphB2 activation, we detected phosphorylation on Y604 which occurs upon receptor clustering following interaction with ephrinB. We found that there is a major increase in Y604 phosphorylation of EphB2 by 5 min of EphB2-ephrinB1 cell contact, which has declined by 15 min, and returned to baseline levels by 30 min (Fig.1B). The transient nature of EphB2 activation likely reflects that most EphB2 is removed from the cell surface by endocytosis during the duration of these experiments and consequently is not available for further activation (Wu et al., 2019). Detection of Y1135 phosphorylation of SHIP2 revealed that there is a similar time course of EphB2 and SHIP2 phosphorylation (Fig.1B). To determine whether SHIP2 phosphorylation requires the kinase activity of EphB2, we carried out analogous experiments with cells expressing kinase-inactive EphB2 (kiEphB2) (Taylor et al., 2017) and found no increase in SHIP2 Y1135 phosphorylation (Fig.1B).

Previous studies have shown that the SAM domain of EphA2 interacts with the SAM domain of SHIP2, but the SAM domain of EphB2 cannot bind to SHIP2 (Lee et al., 2012; Wang et al., 2018; Zhuang et al., 2007). However, it remained possible that EphB2 and SHIP2 do interact, but through some other domain. Indeed, in pull-down experiments with EphB2 and SHIP2 we found that they co-immunoprecipitate (Fig.2A). Consistent with the previous work (Lee et al., 2012; Wang et al., 2018; Zhuang et al., 2007), this binding still occurs for a deletion mutant of SHIP2 lacking the SAM domain (Fig.2B). A potential alternative mode of interaction is through the SH2 domain of SHIP2. To test this, we expressed a fusion protein of GFP and full length SHIP2, or GFP and the SH2 domain of SHIP2, and carried out pull down experiments with EphB2. We found that both fusion proteins co-immunoprecipitate with EphB2, suggesting that they interact through the SH2 domain of SHIP2 (Fig.2C). Since SH2 domains bind to phosphorylated tyrosine motifs, the interaction is likely due to the low level basal activation of EphB2 by endogenous ephrins (Taylor et al., 2017). We therefore tested whether the interaction between EphB2 and SHIP2 changed upon activation of EphB2 by ephrinB1 cells and found that it greatly increased the binding (Fig.2D). Taken together, these findings reveal that EphB2 activation leads to recruitment of SHIP2 through binding to its SH2 domain and the phosphorylation of SHIP2 on Y1135.

**Figure 2.**
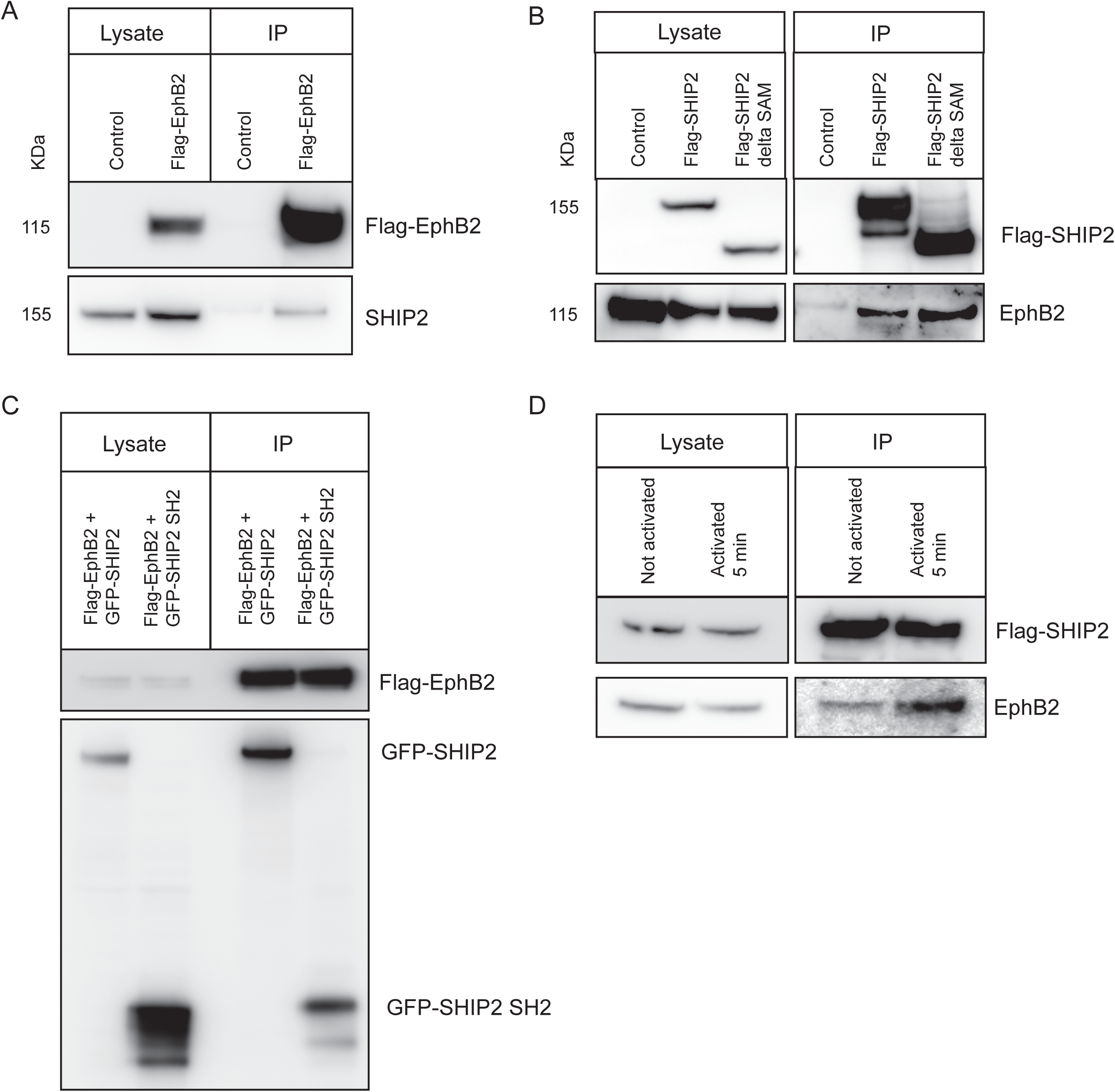
Interaction between EphB2 and SHIP2. Experiments were carried out to analyse the interaction between EphB2 and SHIP2. (A): HEK293 cells were transiently transfected with N-terminally tagged Flag-EphB2 construct. Flag-EphB2 was immunoprecipitated from the lysates and the precipitates analysed by western blotting. SHIP2 was found to co-immunoprecipitate with Flag-EphB2. (B): To determine if the interaction was driven by the SAM domain of SHIP2, Flag-SHIP2 (full length) and Flag-SHIP2 1-847 (delta SAM) was transiently transfected into EphB2 cells. Flag-SHIP2 (full length and delta SAM) was immunoprecipitated from the lysates. EphB2 was found to co-immunoprecipitate with both full length and delta SAM SHIP2. (C): To determine if the interaction is dependent on SH2 domain interactions, Flag-EphB2 and GFP-SHIP2 (full length) or GFP-SHIP2 1-120 were transiently co-transfected in HEK293 cells. Flag-EphB2 was immunoprecipitated from the lysates. The SH2 domain of SHIP2 is sufficient for the interaction with EphB2. (D): To test if the interaction is strengthened by EphB2 activation, Flag-SHIP2 was transiently expressed in EphB2 cells. The EphB2 cells were then mixed with either HEK293 cells (not activated) or mixed with ephrinB1 cells (activated). The cells were briefly centrifuged to force the cells into contact and incubated for 5 min before lysis. Flag-SHIP2 was immunoprecipitated from the lysates. Activation of EphB2 increased the interaction with SHIP2.

### SHIP2 contributes to segregation of EphB2 and ephrinB1 cells

Since the phosphorylation of SHIP2 on Y1135 increases its phosphatase activity (Prasad et al., 2009), our findings raise the question of whether SHIP2 contributes to cell responses to EphB2 activation. A large-scale screen of phosphorylation targets found that SHIP2 knockdown decreases the segregation of EphB2 and ephrinB1 cells (Jorgensen et al., 2009), but the specific nature of the altered segregation or changes in cell responses were not addressed in this study. In previous work, we have shown that cell segregation is mediated by cell repulsion that occurs bi-directionally in EphB2 and ephrinB1 cells (Taylor et al., 2017; Wu et al., 2019). Although SHIP2 phosphorylation is not altered by reverse signaling, the basal level of SHIP2 activity could contribute to responses to ephrinB1 activation. We therefore carried out segregation assays with EphB2 and ephrinB1 cells after SHIP2 knockdown in EphB2 cells, in ephrinB1 cells, or in both cell populations. In these assays, EphB2 and ephrinB1 cells are mixed and plated at sub-confluent density, then incubated for 48 h. In control assays, EphB2 cells segregate to form clusters surrounded by ephrinB1 cells (Fig.3A). We found that segregation still occurs after knockdown of SHIP2 in EphB2 cells, but the EphB2 cell clusters are smaller than in control assays (Fig.3B; quantitated in Fig.3I). Unexpectedly, knockdown of SHIP2 in ephrinB1 cells led to a change in segregation, in which the EphB2 cell clusters are larger (Fig.3C, I) compared with control assays. Knockdown of SHIP2 in both cell populations led to segregation to form clusters similar in size to EphB2/ephrinB1 cell controls (Fig.3D, I).

**Figure 3.**
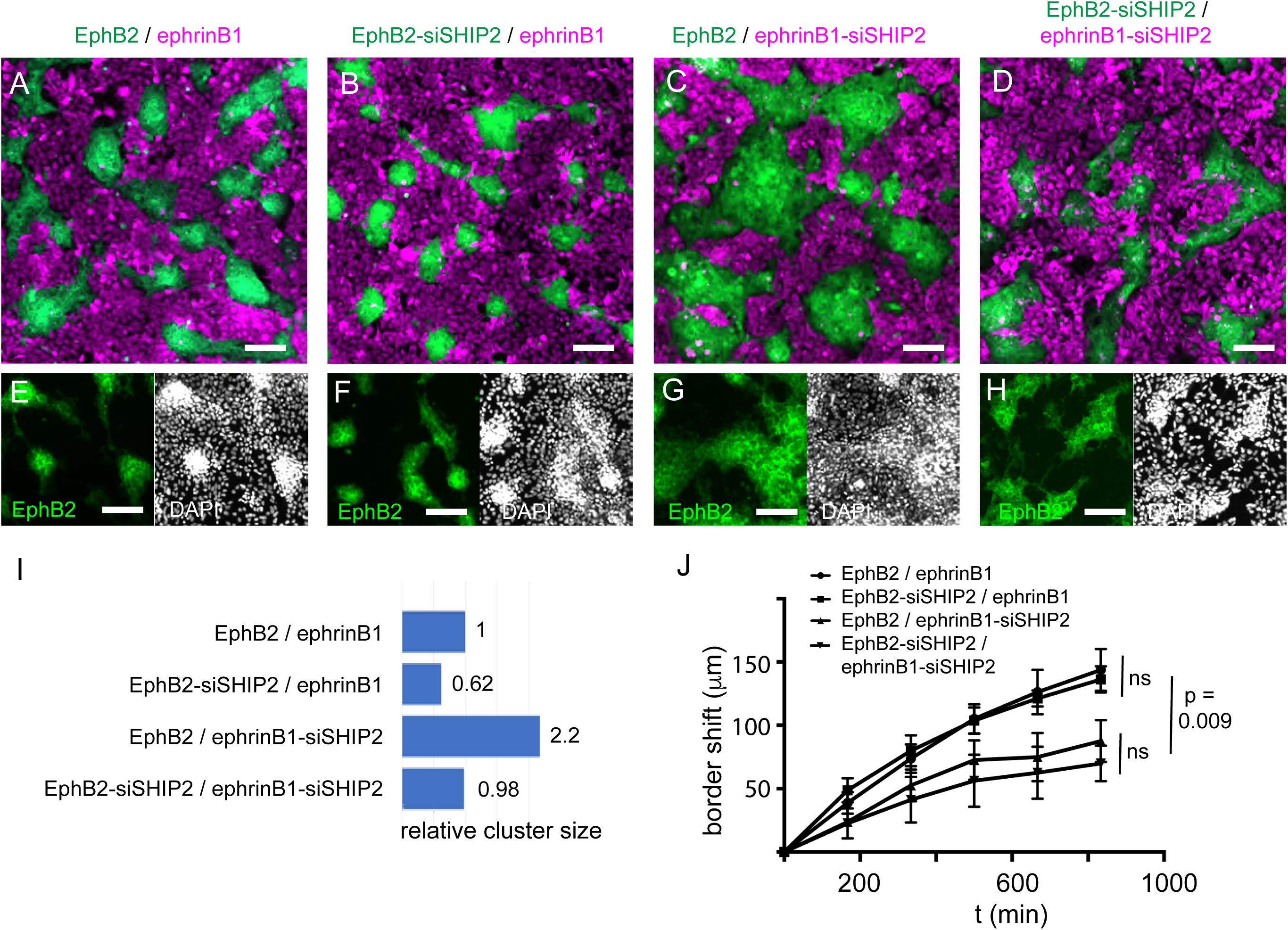
Effect of SHIP2 knockdown on cell segregation. (A-H): Cell segregation assays were carried out in which EphB2 and ephrinB1 cells are mixed and plated at sub-confluent density on tissue culture dishes, then incubated for 48 h. EphB2 cells (green) and ephrinB1 cells (magenta) were labelled with cell tracker dyes, and colours changed using Fiji. (A) EphB2 / ephrinB1 control; (B) SHIP2 knockdown in EphB2 cells; (C) SHIP2 knockdown in ephrinB1 cells; (D) SHIP2 knockdown in both cell populations. (E-H): same assays as shown in the corresponding panel above, but with DAPI staining to visualise the compaction that occurs in EphB2 cell clusters. (I): quantitation of average EphB2 cell cluster size determined by thresholding and particle analysis in Fiji, normalised to control EphB2 / ephrinB1 assay. (J) Quantitation of border shift in boundary assays. Snapshots are shown in Suppl. Fig.1. Scale bars: 100 μm.

We observed that in some combinations of SHIP2 knockdown, there is a decrease in the density of the clustered EphB2 cells after segregation, which can be visualised by DAPI staining (Fig.3E-H). In control EphB2/ephrinB1 cell assays, the EphB2 cells become compacted and pile on top of each other due to the heterotypic repulsion by ephrinB1 cells (Taylor et al., 2017). We found that compaction still occurred in EphB2-siSHIP2 cells (Fig.3F), but was reduced after knockdown of SHIP2 in ephrinB1 cells (Fig.3G, H). The compaction is due to movement of the EphB2-ephrinB1 border, which can be quantitated in boundary assays in which EphB2 cells and ephrinB1-expressing cells are plated either side of a barrier which is then removed: when the two populations meet, the border shifts away from the ephrin-expressing cells (Porazinski et al., 2016; Taylor et al., 2017)(Suppl. Fig.1). We found that following knockdown of SHIP2 in EphB2 cells, the movement of the EphB2-ephrinB1 border was not significantly different from controls (Suppl. Fig.1; quantitated in Fig.3J). Consistent with the findings from segregation assays, the border shift significantly decreased after knockdown of SHIP2 in ephrinB1 cells, and also following knockdown in both cell populations (Suppl. Fig.1; Fig.3J).

### SHIP2 knockdown decreases cell repulsion responses of EphB2 and ephrinB1 cells

These findings raise the question of what change in cell responses to EphB2 and ephrinB1 activation underlie the altered segregation and compaction of EphB2 cells following SHIP2 knockdown. To address this, we made time lapse movies of low density cultures of EphB2 and ephrinB1 cells. As shown previously (Taylor et al., 2017), both EphB2 and ephrinB1 cells have a repulsion response following heterotypic contact, in which there is a local collapse of processes to form a concave surface, concommitant with movement away from the other cell (Fig.4A-A”; Movie 1). In addition, repulsion occurs following homotypic contacts due to low level endogenous expression of ligand, and this is weaker than heterotypic repulsion (Taylor et al., 2017). Knockdown of SHIP2 in EphB2 cells leads to a major change in behaviour, in which they extend dynamic processes in all directions and have a significant decrease in collapse response following heterotypic interactions (p = 0.0002), whereas ephrinB1 cells are still repelled (Fig.4B-B”; Movie 2; quantitated in Fig.4D). In addition, the EphB2-siSHIP2 cells were seen to have reduced homotypic repulsion and to aggregate with each other. These findings suggest that SHIP2 is required for the repulsion response of EphB2 cells.

We found that there is also a major change in the behaviour of ephrinB1-siSHIP2 cells, with a significant decrease in collapse responses (p < 0.0001) following heterotypic interactions, but EphB2 cells are still repelled (Fig.4C-C”, D; Movie 3). As for EphB2-siSHIP2 cells, they have decreased homotypic repulsion and aggregate with each other. Notably, ephrinB1-siSHIP2 cells have greatly reduced motility and a rounded morphology, but unlike EphB2-siSHIP2 cells they do not have rapid cycles of extension and withdrawal of cell processes. Thus, SHIP2 is required for a repulsion response of ephrinB1 cells.

**Figure 4.**
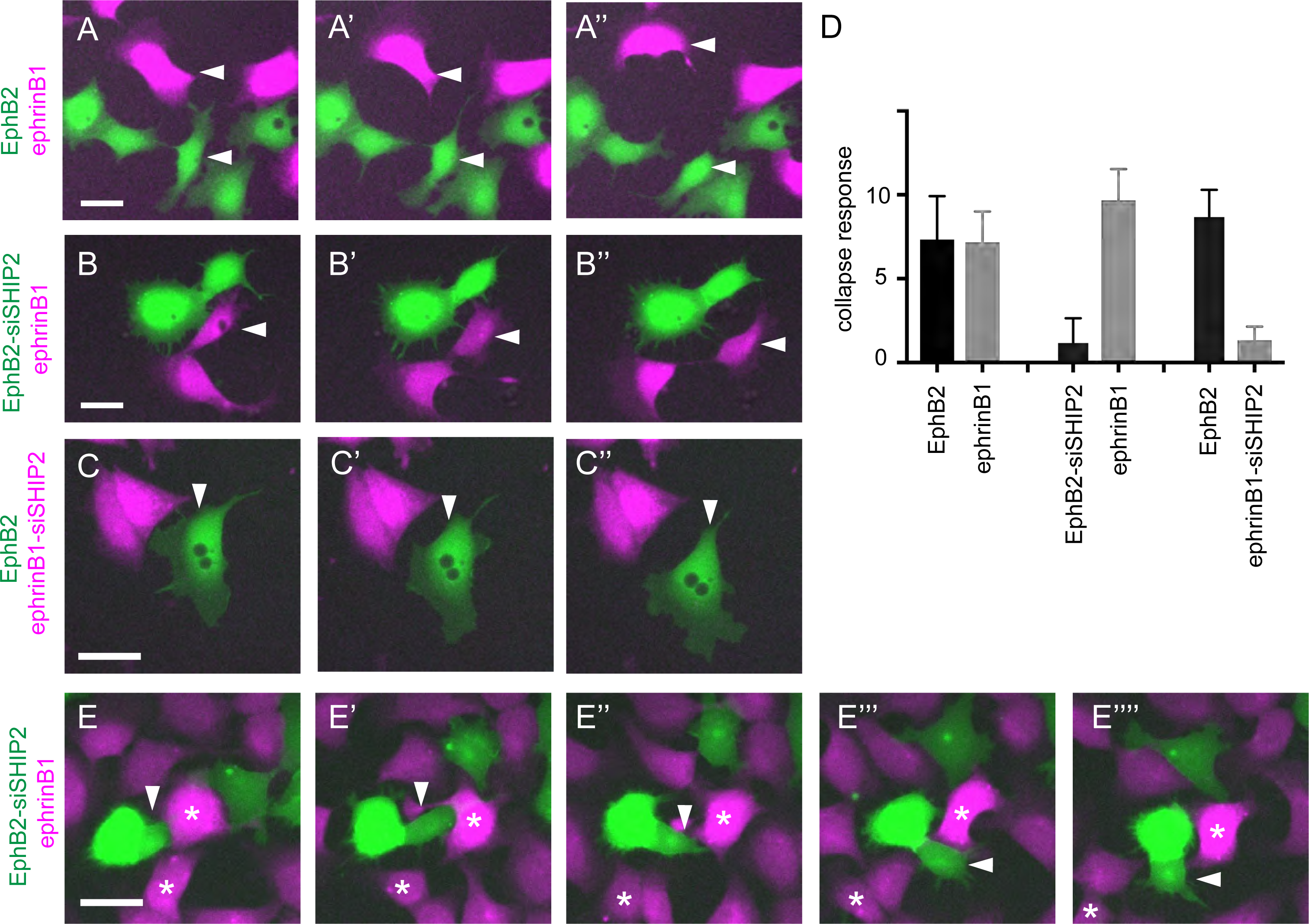
Effect of SHIP2 knockdown on cell repulsion. (A-C): Snapshots of time lapse movies (Movies 1-3, respectively) to illustrate cell repulsion response following interaction of EphB2 (green) and ephrinB1 (magenta) cells, with or without SHIP2 knockdown, indicated with arrowheads. Cell repulsion is characterised by a change of shape due to collapse of cell processes, and movement away from the other cell. (A-A”) EphB2 and ephrinB1 cells are both repelled following their interaction; (B-B”) EphB2-siSHIP2 cell has multiple cell processes and is not repelled, whereas ephrinB1 cell is repelled following their interaction; (C-C”) ephrinB1-siSHIP2 cell is not repelled, whereas EphB2 cell is repelled following their interaction. (D): quantitation of collapse responses from manual tracking of cells in time lapse movies (Movies 4, 5, 7). (E): Snapshots of time lapse movie (Movie 6) showing example of intercalation of EphB2-siSHIP2 cell (arrowhead) between ephrinB1 cells (asterisks). Scale bars: 20 μm.

To gain insights into how the altered cell behaviour leads to changes in cell segregation, we made time lapse movies at late stages when cells are at high density. In control EphB2/ephrinB1 assays, both cell populations are highly motile, have heterotypic repulsion responses, and repeatedly contact each other during segregation (Movie 4). As seen at low density, EphB2-siSHIP2 cells do not have a collapse response upon interacting with ephrinB1 cells (Movie 5). EphB2-siSHIP2 cells are seen to intercalate between ephrinB1 cells, indicative of a decrease in forward repulsion while ephrinB1 cells are still repelled (example shown in Fig.4E-E”; Movie 6). However, most EphB2-siSHIP2 cells aggregate to form stable clusters due to the decrease in homotypic repulsion (Movie 5). The segregation that occurs in EphB2-siSHIP2/ephrinB1 assays is therefore likely driven by heterotypic repulsion of ephrinB1 cells, perhaps in combination with increased homotypic aggregation of EphB2 cells. Smaller EphB2 cell clusters are formed compared with control assays, and this may reflect that heterotypic repulsion of ephrinB1 cells is weaker and a less strong driver of segregation than heterotypic repulsion of EphB2 cells (Taylor et al., 2017; Wu et al., 2019).

Time lapse movies of EphB2/ephrinB1-siSHIP2 cells at high density revealed that ephrinB1-siSHIP2 cells have greatly reduced motility and aggregate with each other, while EphB2 cells have heterotypic repulsion responses (Movie 7). The extensive segregation is therefore likely driven by the strong heterotypic repulsion of EphB2 cells, perhaps in combination with increased homotypic aggregation of ephrinB1 cells. In EphB2/ephrinB1 segregation assays, the highly motile cells can repeatedly contact each other, leading to repeated repulsion of EphB2 cells that pushes them together (Movie 4). In contrast, due to their decreased movement, ephrinB1-siSHIP2 cells rarely move into the gap formed following repulsion of EphB2 cells, and consequently there is less frequent heterotypic contact (Movie 7). This suggests that the decreased motility of ephrinB1-siSHIP2 cells may underlie the decrease in compaction of EphB2 cell clusters.

### Analysis of EphB2 endocytosis

Previous studies have suggested that SHIP2 can inhibit clathrin-mediated endocytosis (Nakatsu et al., 2010; Zhuang et al., 2007) by dephosphorylating phosphoinositides that promote endocytosis (Posor et al., 2015), whereas it promotes endophilin-mediated endocytosis (Boucrot et al., 2015). EphB-ephrinB interactions can lead to endocytosis of the complex in the forward and/or reverse direction, and this influences the strength of cell repulsion responses (Evergren et al., 2018; Gaitanos et al., 2016; Gong et al., 2019; Marston et al., 2003; Pitulescu and Adams, 2010; Zimmer et al., 2003). Furthermore, SHIP2 binds to EphA2 and inhibits its forward endocytosis (Zhuang et al., 2007). It was therefore possible that SHIP2 knockdown changes cell behaviour in part by increasing or decreasing endocytosis. To address this, we carried out knockdowns of SHIP2 and analysed the subcellular distribution of EphB2 and phospho-Y604 EphB2 shortly after contact of EphB2 and ephrinB1 cells. We found that in EphB2/ephrinB1 controls, phospho-Y604 EphB2 is located in EphB2 cells, often in association with retraction fibres, or in intracellular vesicles, but is only rarely found in the interacting ephrinB1 cells (Fig.5A). Furthermore, phospho-Y604 EphB2 co-localises with Rab11, a marker of intracellular vesicles (Fig.5A’). Endocytosis thus occurs mainly in the forward direction following interaction of EphB2 and ephrinB1 cells. No significant difference in the direction of endocytosis was detected after knockdown of SHIP2 in EphB2 cells or ephrinB1 cells (Fig.5B, B’, C, C’). To further address whether SHIP2 knockdown affects the direction of endocytosis, we plated EphB2 and ephrinB1 cells at non-confluent density and allowed them to undergo multiple interactions for 4 h, by which time most EphB2 has been endocytosed. We found that nearly all endocytosis occurs in the forward direction, with EphB2 detected in only few puncta in ephrinB1 cells in controls or after SHIP2 knockdown in EphB2 cells or ephrinB1 cells (Suppl. Fig.1A-C). It remained possible that SHIP2 knockdown has an effect on the timing of EphB2 endocytosis that is not detected in these assays. Since endocytosis terminates further activation of EphB2, indirect evidence might come from analysis of EphB2 phosphorylation. We therefore used the EphB2/ephrinB1 cell mixing assay described above (Fig.1B) for time course analyses of EphB2 phosphorylation following knockdown of SHIP2 in EphB2 or ephrinB1 cells. We found that knockdown of SHIP2 in EphB2 cells led to no change in the time course of EphB2 phosphorylation compared with controls (Fig.5D; quantitated in Fig.5E). In contrast, knockdown of SHIP2 in ephrinB1 cells led to an increase in EphB2 phosphorylation at 5 and 15 min of activation (Fig.D, E). This increase in EphB2 phosphorylation may underlie the increase in collapse response of EphB2 cells when interacting with ephrinB1-siSHIP2 cells compared with ephrinB1 cells (Fig.4D).

**Figure 5.**
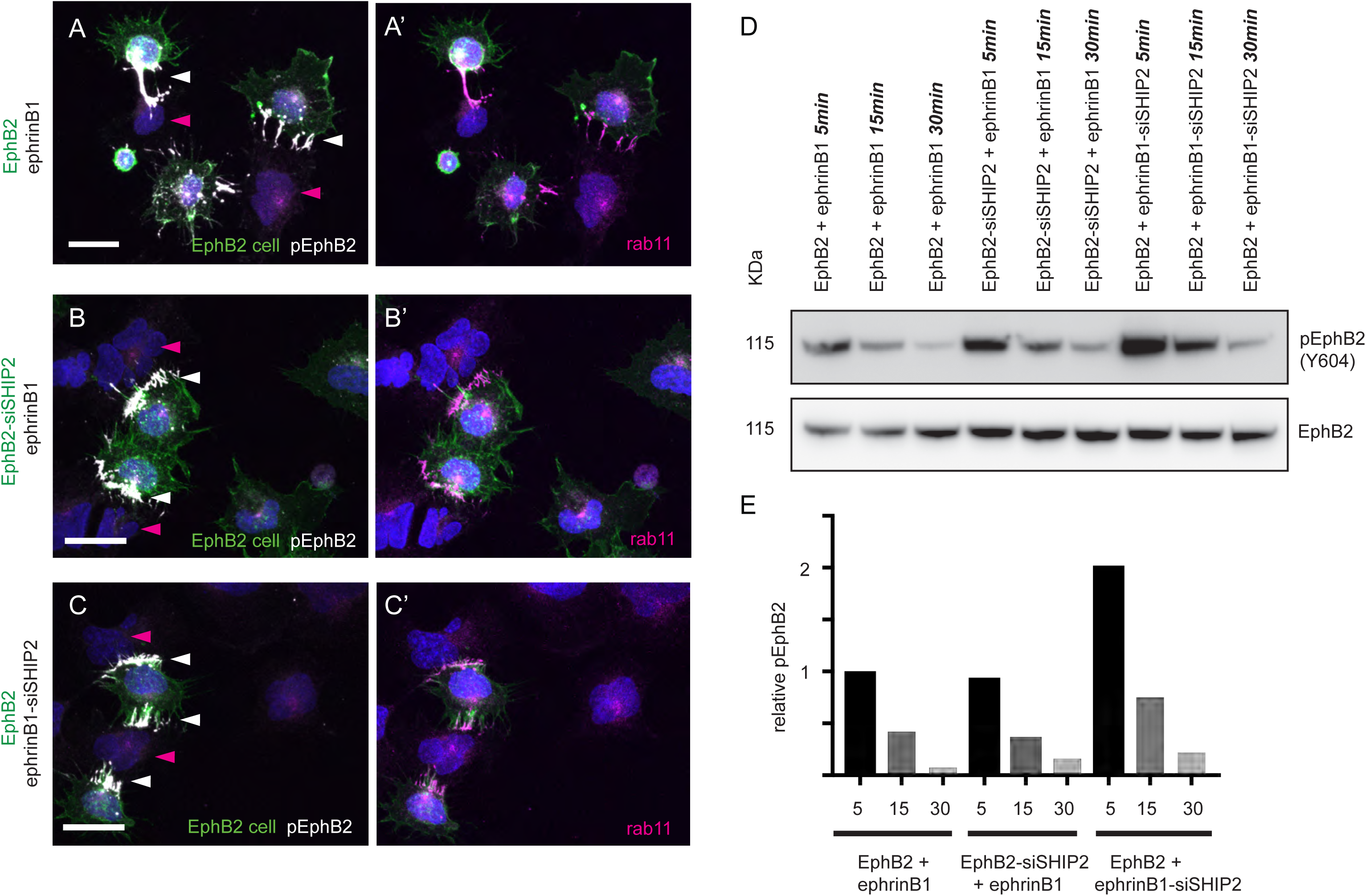
Analysis of endocytosis and activation of EphB2 following SHIP2 knockdown. (A-C): immunostaining to detect pEphB2 (Y604) (white signal) and rab11, which marks intracellular vesicles (magenta signal), after 15 min of interaction. (A-A’) control EphB/ephrinB1 assay; (B-B’) after SHIP2 knockdown in EphB2 cells; (C-C’) after SHIP2 knockdown in ephrinB1 cells. EphB2 cells are labelled in green (examples indicated with white arrowheads) and ephrinB1 cells are unlabelled (magenta arrowheads). pEphB2 and rab11 are detected in retraction fibres and intracellular vesicles of EphB2 cells. Scale bars: 20 μm. (D) Western blot analysis of EphB2 phosphorylation (Y604) after forced interaction of EphB2 and ephrinB1 cells, with or without SHIP2 knockdown. (E) Quantitation of pEphB2 relative to EphB2 / ephrinB1 control at 5 min. There is no change in EphB2 phosphorylation following knockdown of SHIP2 in EphB2 cells, but it increases at 5 and 15 min following knockdown in ephrinB1 cells. This increase was found in two independent experiments, and the average is shown in (E).

## DISCUSSION

A proteomic analysis of changes in tyrosine phosphorylation downstream of EphB2 activation found SHIP2 as one of the targets, which together with a knockdown screen suggested a role in EphB2/ephrinB1 cell segregation (Jorgensen et al., 2009). One potential mechanism of SHIP2 action is suggested by studies showing that it interacts with the SAM domain of EphA2 and inhibits endocytosis (Zhuang et al., 2007). Other studies have implicated SHIP2 in the polarisation and migration of cells (Elong Edimo et al., 2016; Kato et al., 2012; Lam et al., 2012; Venkatareddy et al., 2011; Yu et al., 2010), although this has not been studied in the context of cell responses to Eph receptor signaling. Here, we show that SHIP2 binds to EphB2, but through a different mode of interaction from the binding to EphA2. In addition, we show that SHIP2 is required for cell repulsion responses, both of EphB2 and ephrinB1 cells, that are implicated in cell segregation.

### Binding of SHIP2 to EphB2

Previous work has shown that SHIP2 binds to EphA2 through interactions between their SAM domains, and that such binding does not occur for the SAM domain of EphB2 (Lee et al., 2012; Wang et al., 2018; Zhuang et al., 2007). We found that SHIP2 interacts with EphB2, and does so through the SH2 domain of SHIP2. Since SH2 domains bind to phosphotyrosine motifs, this suggests that SHIP2 is recruited to activated EphB2, and indeed we found increased binding of SHIP2 upon activation by ephrinB1. We confirmed the results of phosphoproteomic studies (Jorgensen et al., 2009) showing that activation of EphB2 leads to increased phosphorylation of SHIP2 at Y1135, whereas this does not occur during reverse signaling through ephrinB1. Since phosphorylation of SHIP2 at Y1135 increases its phosphatase activity (Prasad et al., 2009), this suggests that activation of EphB2 leads to recruitment and increased activity of SHIP2 at the site of heterotypic contact of EphB2 and ephrinB1 cells. This raises the question of what role SHIP2 may play in cell responses to EphB2 activation.

### Effect of SHIP2 knockdown on cell segregation and compaction

Analysis of SHIP2 function in EphB2/ephrinB1 cell segregation assays found that knockdown in EphB2 cells leads to a decreased size of EphB2 cell clusters, suggesting a partial disruption of segregation. In contrast, knockdown of SHIP2 in ephrinB1 cells did not disrupt segregation, and surprisingly leads to an increase in the size of EphB2 cell clusters. These findings can be explained by changes in the responses of EphB2 and ephrinB1 cells that have distinct contributions to cell segregation. Previous work has shown that the strong cell repulsion response to EphB2 activation has the dominant role in segregation (O’Neill et al., 2016; Rohani et al., 2011; Rohani et al., 2014; Taylor et al., 2017), but there is some contribution from the weaker repulsion that occurs in ephrinB1 cells (Wu et al., 2019). We found that SHIP2 knockdown decreased heterotypic cell repulsion responses both in EphB2 cells and in ephrinB1 cells. In addition, SHIP2 knockdown led to homotypic cell aggregation, concommitant with a decrease in the cell repulsion that can occur following homotypic interactions, which is due to low level endogenous expression of ligands (Taylor et al., 2017). Taken together, these observations suggest that in EphB2-siSHIP2/ephrinB1 assays, there is less segregation because forward repulsion is decreased. The remaining segregation may be driven by reverse repulsion and/or residual levels of forward repulsion. In favour of the latter idea, there is a major decrease in cluster size in EphB2-siSHIP2 / ephrinB1-siSHIP2 assays compared with EphB2 / ephrinB1-siSHIP2 assays, in which reverse repulsion is decreased in both assays. Furthermore, measurements and simulations suggest that even a small difference between heterotypic and homotypic forward repulsion or tension can drive segregation (Canty et al., 2017; Taylor et al., 2017). In contrast, forward repulsion is intact and likely drives the extensive segregation that occurs in EphB2/ephrinB1-siSHIP2 assays. However, this does not account for the increased size and lower compaction of EphB2 cell clusters in these latter assays.

Previous work found that when Eph- and ephrin-expressing cell populations are juxtaposed in cell culture assays, the repulsion response of Eph cells causes the border to move away from the ephrin cells (Porazinski et al., 2016; Taylor et al., 2017). When clusters of Eph-expressing cells are surrounded by ephrin-expressing cells, this border shift compacts the Eph cell clusters (Taylor et al., 2017) and can cause extrusion of cells from epithelia (Porazinski et al., 2016). We found that in assays with EphB2 and ephrinB1-siSHIP2 cells, the border shift and compaction of EphB2 cells is decreased compared with control EphB2/ephrinB1 cell assays. Analysis of time lapse movies at low cell density suggests that the decrease is due to the lower motility of ephrinB1-siSHIP2 cells, which reduces the frequency of them meeting again the EphB2 cells that have moved away. Likewise, in boundary assays, there is slower movement of the front of migrating ephrinB1-siSHIP2 cells. Based on these observations, we propose that for migratory cells the compaction of EphB2 cell clusters is affected by the motility of ephrinB1 cells.

### Potential role of SHIP2 in endocytosis

Since previous studies have shown that SHIP2 inhibits the endocytosis of EphA2 (Zhuang et al., 2007), we wondered whether it may have a similar function for EphB2. Although we did not observe that SHIP2 knockdown increases endocytosis of EphB2, this may be because endocytosis is rapid and nearly all occurs in the forward direction following interaction of EphB2 and ephrinB1 cells. There is also a technical difference, since the EphA2 study (Zhuang et al., 2007) used clustered soluble ephrins to activate receptor, and endocytosis of Eph receptor has been found to involve some different players when activated by membrane-bound or soluble ephrins (Gaitanos et al., 2016). We also tested the possibility that SHIP2 acts in ephrinB1 cells to inhibit reverse endocytosis. Although we did not detect a significant increase in reverse endocytosis after SHIP2 knockdown in ephrinB1 cells, this could reflect that other factors are sufficient for the strong bias towards forward endocytosis of EphB2-ephrinB1 complexes. Intriguingly, in time course analysis of activation following forced interaction of EphB2 and ephrinB1 cells, we found that knockdown of SHIP2 in ephrinB1 cells increases the initial level of EphB2 phosphorylation. One possible explanation of this non-autonomous effect is that SHIP2 knockdown in ephrinB1 cells alters the balance of endocytic mechanisms, which delays rather than blocks forward endocytosis of EphB2, and thus increases its activation by ephrinB1 at the cell surface. Further studies will be required to definitively address whether SHIP2 contributes to regulation of the endocytosis of EphB2 and/or ephrinB1.

### SHIP2 in cell repulsion

Previous studies have shown that SHIP2 has important roles in the polarisation of cells required for migration, by dephosphorylating phosphoinositides to affect the balance and localisation of PI(3,4,5)P3, PI(3,4)P3 and PI(4,5)P2 that regulate actin filament formation (Charest and Firtel, 2006; Krause and Gautreau, 2014). In some cell types, SHIP2 knockdown leads to an increase in migration (Lam et al., 2012), which for N1 glioblastoma cells is because PI(4,5)P2 promotes migration and is degraded by SHIP2 (Elong Edimo et al., 2016). However, in other cell types SHIP2 is required for polarisation and directional migration by dephosphorylating PI(3,4,5)P3 to generate PI(3,4)P2 (Krause and Gautreau, 2014) and also through interactions with RhoA (Kato et al., 2012). This latter situation is found for EphB2 and ephrinB1 cells, in which SHIP2 knockdown leads to a decrease in cell repulsion responses which require localised collapse of the actin cytoskeleton. Since SHIP2 is recruited to activated EphB2, and its activity is increased by phosphorylation on Y1135, cell repulsion may involve an increase in PI(3,4)P2 levels at the site of EphB2 activation. In the case of ephrinB1 cells, there is no recruitment or increased phosphorylation of SHIP2, but its constitutive activity may be required for correct intracellular polarisation of phosphoinositides. It will be interesting to determine whether this different relationship with SHIP2 contributes to the more persistent migration of EphB2 cells compared with ephrinB1 cells after heterotypic repulsion (Taylor et al., 2017). It will also be important to address whether there is an interplay with PI3-kinase, as this has been implicated in cell responses to Eph receptor activation (Hunter et al., 2006; Kaplan et al., 2012; Lin et al., 2015; Pasquale, 2008; Steinle et al., 2002) and is a key enzyme for phosphoinositide metabolism by generating PI(3,4,5)P3 from PI(4,5)P2.

## Acknowledgements

We thank the staff of Cell Services and of the Light Microscopy Technology Platform at the Francis Crick Institute for their invaluable support. This work was supported by the Francis Crick Institute which receives its core funding from Cancer Research UK (FC001217), the UK Medical Research Council (FC001217), and the Wellcome Trust (FC001217).

## Author contributions

Tim Ashlin: Conceptualisation, Investigation, Methodology, Writing – review and editing.

Zhonglin Wu: Investigation, Writing – review and editing.

Qiling Xu: Investigation, Writing – review and editing.

David Wilkinson: Conceptualisation, Formal analysis, Funding acquisition; Writing – original draft, Supervision.

## MATERIALS AND METHODS

### Cell culture assays

EphB2-expressing and ephrinB1-expressing HEK293 cell lines (Poliakov et al., 2008) were used for cell culture assays for cell segregation and border formation, as described in more detail previously (Taylor et al., 2017; Wu et al., 2019). Cells were labelled with CMFDA (green) or CMRA (red) cell tracker dyes (Molecular Probes, Invitrogen). In segregation assays, EphB2 and ephrinB1 cells were mixed in equal proportions, plated on a fibronectin-coated slide (Lab-Tek) at a density of 200,000 cell/cm^2^, and cultured for 48 h. To visualise cell responses, 20,000 cells were placed into a 0.7 cm^2^ well and images were taken every 3 min. Time lapse images were processed using ImageJ, with red colour changed to magenta so that the colours can be more easily distinguished. In boundary assays, a two well insert (Ibidi) was placed onto a fibronectin-coated chambered slide and 88 μl of cells at 1 × 10^6^ cells/ml put into each side. Cells were incubated for 4 h before lifting the barrier, and then incubated for a further 40 h. Images were taken at 20 min intervals.

### Activation of EphB2 by cell mixing

The assay to synchronously activate EphB2 has been described previously (Wu et al., 2019). In brief, EphB2, kiEphB2 and ephrinB1 cell lines were harvested separately and adjusted to 6 × 10^6^ cells/ml. 0.25 ml of each were then mixed in a 1.5 ml tube, and centrifuged at 180 rpm for 45 sec to force the cells into contact. The pellets were incubated at 37°C for the appropriate time, and then cells lysed with RIPA buffer (50 mM Tris/HCl pH 8, 150 mM NaCl, 1% Triton-X100, 0.5% Sodium Deoxycholate, 0.1% SDS) supplemented with protease inhibitors (Sigma Aldrich) on ice. The lysates were briefly sonicated and then cleared by centrifugation at 13,300 rpm for 15 min at 4°C. The BCA assay was used to determine the protein concentration of samples. Antibodies used for Western blot analysis were to detect: EphB2 and GAPDH from Thermo Fisher (371700, MA5-15738, respectively); phosphorylated EphB2 from Abcam (ab61791); ephrinB1/2/3 from Santa Cruz (sc-910); FLAG tag from Sigma Aldrich (F3165); SHIP2 and phosphorylated SHIP2-Y1135 from Cell Signaling Technologies (#2839 and 5445, respectively).

### Activation with Fc reagents

1.5 × 10^6^ EphB2 or ephrinB1 cells were seeded onto a 10 cm dish and cultured overnight to allow the cells to adhere. The media was replaced with 3 ml of complete DMEM media. The cells were then stimulated with 1 μg/ml of the appropriate Fc-conjugates. Fc-conjugates for mouse EphB2 (467-B2-200) and mouse ephrinB1 (473-EB-200) were obtained from R&D Systems. Following the incubation period, the cells were lysed with RIPA buffer supplemented with protease inhibitors on ice, briefly sonicated and then cleared, as described above.

### Immunoprecipitations

Cells were harvested in NP-40 buffer (20 mM Tris-HCl pH 8, 137 mM NaCl, 1% NP-40 and 2 mM EDTA) supplemented with protease and phosphatase I and II inhibitors (Sigma–Aldrich, P8340, P2850 and P5726 respectively). The cells were sonicated, the lysates were cleared by centrifugation, and protein concentration determined as described above. For immunoprecipitation, 1 mg protein was incubated with 50 μl equilibrated anti-FLAG M2 affinity gel (Sigma–Aldrich, A2220) on a rotating wheel for 1 h at 4 °C. The beads were washed four times with NP-40 buffer; the final wash was carried out in a fresh Eppendorf tube. After the final centrifugation, the supernatant was aspirated and replaced with 30 μl 2X NuPAGE LDS sample buffer under reducing conditions (Life Technologies).

### Immunocytochemistry

Cells were fixed for 15 min in 4% paraformaldehyde at 37°C, washed three times in PBS and stored at 4°C prior to immunostaining. Cells were washed twice in PBT (0.1% Tween20 in PBS), blocked for 30 min and then incubated for 2-3 h with primary antibodies in 2.5% goat serum, 1% DMSO in PBT. After washing eight times in PBT during 1 h, cells were incubated for 2 h in secondary antibody (donkey anti-mouse Cy5-conjugated and donkey anti-rabbit Cy3-conjugated, Jackson ImmunoResearch, 1:400) together with DAPI nuclear counterstain. The cells were then washed eight times in PBT and mounted in FluorSave. Immunostained cells were imaged using a Zeiss LSM700 confocal microscope and images processed using ImageJ. The antibodies used were to detect: EphB2 (see above, 1:300); phospho-EphB2 (see above, 1:300); Rab11 (BD Biosciences; 610656).

### Constructs and transfection

The constructs used to express FLAG-SHIP2 and FLAG-SHIP2 1-847 were obtained through the MRC PPU Reagents and Services facility (mrcppureagents.dundee.ac.uk). The construct used to express FLAG-EphB2 (HG10762-NF) was sourced from Sino Biological Inc. For transfection, 2 × 10^6^ cells were seeded onto a 10 cm dish, cultured overnight, and then the media was replaced with 10 ml of complete DMEM media. The cells were then transfected using FugeneHD (Promega), according to the manufacturer’s instructions. To carry out the transfection of a single dish, 10 μg plasmid DNA was diluted in 500 μl OPTIMEM and 30 μl FugeneHD was added. The resulting solution was incubated for 10 min at room temperature and then added dropwise to the dish containing the cells. The transfected cells were then cultured for 24 h before using in experiments.

### Treatment of cell lines with SHIP2 siRNA

1.5 × 10^5^ cells were seeded into each well of a 6-well plate and cultured overnight, then the media changed to 2 ml DMEM + GlutaMAX™ supplemented with 10% FBS (no penicillin/streptomycin). The cells were then transfected with SHIP2 siRNAs (SI04371668, SI04360475, SI04267228, SI04207280) using Lipofectamine RNAiMAX (1027415, Qiagen), according to the manufacturer’s instructions. To carry out the transfection of a single well, two tubes were created per knockdown condition. Tube 1 contains 150 μl OPTIMEM and 9 μl Lipofectamine RNAIMAX. Tube 2 contains 150 μl OPTIMEM and 1.5 μl of 20 μM siRNA solution. The two tubes are then mixed 1:1 and incubated for 10 min at room temperature. 250 μl was then added dropwise to the well containing the cells and 2 ml of DMEM media + 10% FBS (no penicillin/ streptomycin). The transfected cells were cultured for 48 h before using to assess knockdown and carry out experiments.

---

**Supplementary Figure 1.**
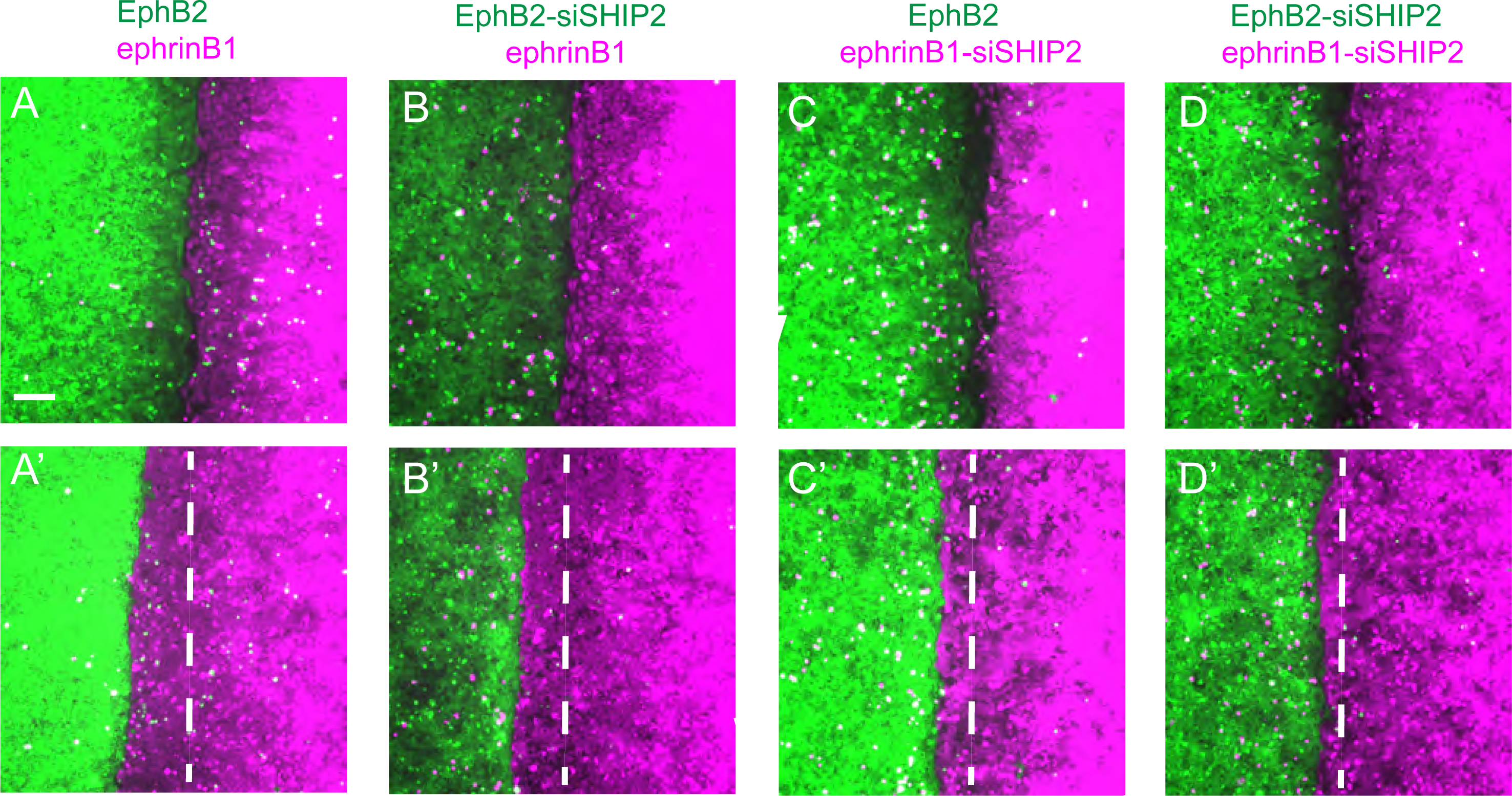
Analysis of SHIP2 knockdown in boundary assays. (A-D): boundary assays were carried out in which EphB2 and ephrinB1 cells meet to form a border. Snapshots are shown of the meeting of the two cell populations (A-D) and at the endpoint of the experiment (A’-D’; original position of border indicated with dashed white line). In control EphB2 / ephrinB1 assays (A, A’) the border shifts away from ephrinB1 cells. The border shift is not affected by knockdown of SHIP2 in EphB2 cells (B, B’) but decreases following knockdown in ephrinB1 cells (C, C’) or both populations (D, D’). The border shift is quantitated in Fig.3J. Scale bar: 100 μm.

**Supplementary Figure 2.**
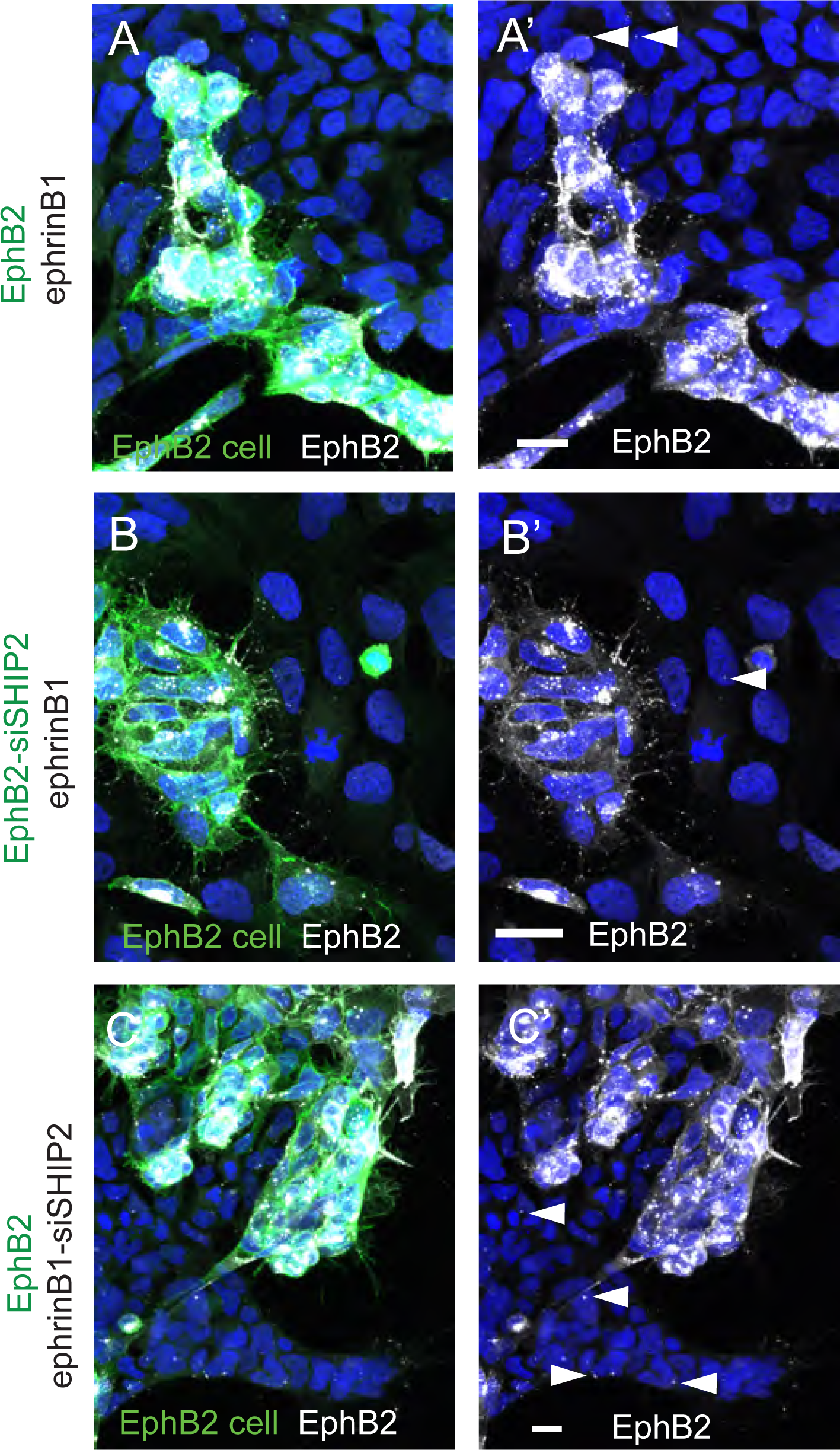
Analysis of endocytosis of EphB2 following SHIP2 knockdown. Immunostaining to detect EphB2 (white signal) after 4 h of plating in segregation assays. (A-A’) control EphB2/ephrinB1 assay; (B-B’) after SHIP2 knockdown in EphB2 cells; (C-C’) after SHIP2 knockdown in ephrinB1 cells. EphB2 cells are labelled in green and ephrinB1 cells are unlabelled. By this late time point, nearly all EphB2 has been endocytosed. Most EphB2 is detected within EphB2 cells, and only a few puncta are present in ephrinB1 cells (examples indicated with arrowheads), suggesting that nearly all endocytosis occurs in the forward direction in all of the conditions tested. Scale bars: 20 μm.

## Movie legends

**Movie 1**

Time lapse movie of EphB2 cells (green) and ephrinB1 cells (magenta). High power view showing repulsion response of both EphB2 and ephrinB1 cells.

**Movie 2**

Time lapse movie of EphB2-siSHIP2 cells (green) and ephrinB1 cells (magenta). Repulsion responses occur for ephrinB1 cells but not for EphB2-siSHIP2 cells.

**Movie 3**

Time lapse movie of EphB2 cells (green) and ephrinB1-siSHIP2 cells (magenta). Repulsion responses occur for EphB2 cells but not for ephrinB1-siSHIP2 cells.

**Movie 4**

Time lapse movie of EphB2 cells (green) and ephrinB1 cells (magenta) at late stage of segregation. Low power view showing repulsion response of both EphB2 and ephrinB1 cells.

**Movie 5**

Time lapse movie of EphB2-siSHIP2 cells (green) and ephrinB1 cells (magenta) at late stage of segregation. Low power view showing repulsion response of ephrinB1 cells and decreased repulsion with increased aggregation of EphB2-siSHIP2 cells.

**Movie 6**

Time lapse movie of EphB2-siSHIP2 cells (green) and ephrinB1 cells (magenta) at late stage of segregation. High power view showing example of intercalation of an EphB2-siSHIP2 cell between ephrinB1 cells.

**Movie 7**

Time lapse movie of EphB2 cells (green) and ephrinB1-siSHIP2 cells (magenta) at late stage of segregation. Low power view showing aggregation and low motility of ephrinB1 cells and repulsion of EphB2 cells.

## References

Astin, J.W., Batson, J., Kadir, S., Charlet, J., Persad, R.A., Gillatt, D., Oxley, J.D., and Nobes, C.D. (2010). Competition amongst Eph receptors regulates contact inhibition of locomotion and invasiveness in prostate cancer cells. Nature Cell Biology 12, 1194–1204.

Batlle, E., and Wilkinson, D.G. (2012). Molecular mechanisms of cell segregation and boundary formation in development and tumorigenesis. Cold Spring Harbor Perspect Biol 4: a008227.

Boucrot, E., Ferreira, A.P., Almeida-Souza, L., Debard, S., Vallis, Y., Howard, G., Bertot, L., Sauvonnet, N., and McMahon, H.T. (2015). Endophilin marks and controls a clathrin-independent endocytic pathway. Nature 517, 460–465.

Calzolari, S., Terriente, J., and Pujades, C. (2014). Cell segregation in the vertebrate hindbrain relies on actomyosin cables located at the interhombomeric boundaries. EMBO J 33, 686–701.

Canty, L., Zarour, E., Kashkooli, L., Francois, P., and Fagotto, F. (2017). Sorting at embryonic boundaries requires high heterotypic interfacial tension. Nat Commun 8, 157.

Cayuso, J., Xu, Q., Addison, M., and Wilkinson, D.G. (2019). Actomyosin regulation by Eph receptor signaling couples boundary cell formation to border sharpness. Elife 8 e49696.

Cayuso, J., Xu, Q., and Wilkinson, D.G. (2015). Mechanisms of boundary formation by Eph receptor and ephrin signaling. Dev Biol 401, 122–131.

Charest, P.G., and Firtel, R.A. (2006). Feedback signaling controls leading-edge formation during chemotaxis. Curr Opin Genetics Development 16, 339–347.

Dahmann, C., Oates, A.C., and Brand, M. (2011). Boundary formation and maintenance in tissue development. Nat Rev Genet 12, 43–55.

Egea, J., and Klein, R. (2007). Bidirectional Eph-ephrin signaling during axon guidance. Trends Cell Biol 17, 230–238.

Elong Edimo, W., Ghosh, S., Derua, R., Janssens, V., Waelkens, E., Vanderwinden, J.M., Robe, P., and Erneux, C. (2016). SHIP2 controls plasma membrane PI(4,5)P2 thereby participating in the control of cell migration in 1321 N1 glioblastoma cells. J Cell Science 129, 1101–1114.

Evergren, E., Cobbe, N., and McMahon, H.T. (2018). Eps15R and clathrin regulate EphB2-mediated cell repulsion. Traffic 19, 44–57.

Fagotto, F. (2014). The cellular basis of tissue separation. Development 141, 3303–3318.

Fagotto, F., Rohani, N., Touret, A.S., and Li, R. (2013). A molecular base for cell sorting at embryonic boundaries: contact inhibition of cadherin adhesion by ephrin/ Eph-dependent contractility. Developmental Cell 27, 72–87.

Fagotto, F., Winklbauer, R., and Rohani, N. (2014). Ephrin-Eph signaling in embryonic tissue separation. Cell Adh Migr 8, 308–326.

Gaitanos, T.N., Koerner, J., and Klein, R. (2016). Tiam-Rac signaling mediates transendocytosis of ephrin receptor EphB2 and is important for cell repulsion. Journal Cell Biology 214, 735–752.

Gale, N.W., Holland, S.J., Valenzuela, D.M., Flenniken, A., Pan, L., Ryan, T.E., Henkemeyer, M., Strebhardt, K., Hirai, H., Wilkinson, D.G., et al. (1996). Eph receptors and ligands comprise two major specificity subclasses and are reciprocally compartmentalized during embryogenesis. Neuron 17, 9–19.

Gong, J., Gaitanos, T.N., Luu, O., Huang, Y., Gaitanos, L., Lindner, J., Winklbauer, R., and Klein, R. (2019). Gulp1 controls Eph/ephrin trogocytosis and is important for cell rearrangements during development. Journal Cell Biology 218, 3455–3471.

Hunter, S.G., Zhuang, G., Brantley-Sieders, D., Swat, W., Cowan, C.W., and Chen, J. (2006). Essential role of Vav family guanine nucleotide exchange factors in EphA receptor-mediated angiogenesis. Mol Cell Biol 26, 4830–4842.

Jorgensen, C., Sherman, A., Chen, G.I., Pasculescu, A., Poliakov, A., Hsiung, M., Larsen, B., Wilkinson, D.G., Linding, R., and Pawson, T. (2009). Cell-specific information processing in segregating populations of Eph receptor ephrin-expressing cells. Science 326, 1502–1509.

Kania, A., and Klein, R. (2016). Mechanisms of ephrin-Eph signalling in development, physiology and disease. Nat Rev Mol Cell Biol 17, 240–256.

Kaplan, N., Fatima, A., Peng, H., Bryar, P.J., Lavker, R.M., and Getsios, S. (2012). EphA2/Ephrin-A1 signaling complexes restrict corneal epithelial cell migration. Invest Ophthalmol Vis Sci 53, 936–945.

Kato, K., Yazawa, T., Taki, K., Mori, K., Wang, S., Nishioka, T., Hamaguchi, T., Itoh, T., Takenawa, T., Kataoka, C., et al. (2012). The inositol 5-phosphatase SHIP2 is an effector of RhoA and is involved in cell polarity and migration. Mol Biol Cell 23, 2593–2604.

Klein, R. (2012). Eph/ephrin signalling during development. Development 139, 4105–4109.

Krause, M., and Gautreau, A. (2014). Steering cell migration: lamellipodium dynamics and the regulation of directional persistence. Nat Rev Mol Cell Biol 15, 577–590.

Lam, P.Y., Yoo, S.K., Green, J.M., and Huttenlocher, A. (2012). The SH2-domain-containing inositol 5-phosphatase (SHIP) limits the motility of neutrophils and their recruitment to wounds in zebrafish. J Cell Science 125, 4973–4978.

Lee, H.J., Hota, P.K., Chugha, P., Guo, H., Miao, H., Zhang, L., Kim, S.J., Stetzik, L., Wang, B.C., and Buck, M. (2012). NMR structure of a heterodimeric SAM:SAM complex: characterization and manipulation of EphA2 binding reveal new cellular functions of SHIP2. Structure 20, 41–55.

Lin, B., Yin, T., Wu, Y.I., Inoue, T., and Levchenko, A. (2015). Interplay between chemotaxis and contact inhibition of locomotion determines exploratory cell migration. Nat Commun 6, 6619.

Marston, D.J., Dickinson, S., and Nobes, C.D. (2003). Rac-dependent trans-endocytosis of ephrinBs regulates Eph-ephrin contact repulsion. Nature Cell Biology 5, 879–888.

Nakatsu, F., Perera, R.M., Lucast, L., Zoncu, R., Domin, J., Gertler, F.B., Toomre, D., and De Camilli, P. (2010). The inositol 5-phosphatase SHIP2 regulates endocytic clathrin-coated pit dynamics. J Cell Biology 190, 307–315.

O’Neill, A.K., Kindberg, A.A., Niethamer, T.K., Larson, A.R., Ho, H.Y.H., Greenberg, M.E., and Bush, J.O. (2016). Unidirectional Eph/ephrin signaling creates a cortical actomyosin differential to drive cell segregation. J Cell Biology 215, 217–229.

Pasquale, E.B. (2008). Eph-ephrin bidirectional signaling in physiology and disease. Cell 133, 38–52.

Pitulescu, M.E., and Adams, R.H. (2010). Eph/ephrin molecules--a hub for signaling and endocytosis. Genes Dev 24, 2480–2492.

Poliakov, A., Cotrina, M.L., Pasini, A., and Wilkinson, D.G. (2008). Regulation of EphB2 activation and cell repulsion by feedback control of the MAPK pathway. J Cell Biology 183, 933–947.

Porazinski, S., de Navascues, J., Yako, Y., Hill, W., Jones, M.R., Maddison, R., Fujita, Y., and Hogan, C. (2016). EphA2 Drives the Segregation of Ras-Transformed Epithelial Cells from Normal Neighbors. Current Biology 26, 3220–3229.

Posor, Y., Eichhorn-Grunig, M., and Haucke, V. (2015). Phosphoinositides in endocytosis. Biochim Biophys Acta 1851, 794–804.

Prasad, N.K., Werner, M.E., and Decker, S.J. (2009). Specific tyrosine phosphorylations mediate signal-dependent stimulation of SHIP2 inositol phosphatase activity, while the SH2 domain confers an inhibitory effect to maintain the basal activity. Biochemistry 48, 6285–6287.

Rohani, N., Canty, L., Luu, O., Fagotto, F., and Winklbauer, R. (2011). EphrinB/EphB Signaling Controls Embryonic Germ Layer Separation by Contact-Induced Cell Detachment. PLoS Biology 9, e1000597.

Rohani, N., Parmeggiani, A., Winklbauer, R., and Fagotto, F. (2014). Variable combinations of specific ephrin ligand/eph receptor pairs control embryonic tissue separation. PLoS Biology 12, e1001955.

Solanas, G., Cortina, C., Sevillano, M., and Batlle, E. (2011). Cleavage of E-cadherin by ADAM10 mediates epithelial cell sorting downstream of EphB signalling. Nature Cell Biology 13, 1100–1107.

Steinberg, M.S. (2007). Differential adhesion in morphogenesis: a modern view. Curr Opin Genetics Development 17, 281–286.

Steinle, J.J., Meininger, C.J., Forough, R., Wu, G., Wu, M.H., and Granger, H.J. (2002). Eph B4 receptor signaling mediates endothelial cell migration and proliferation via the phosphatidylinositol 3-kinase pathway. J Biol Chem 277, 43830–43835.

Tanaka, M., Kamo, T., Ota, S., and Sugimura, H. (2003). Association of Dishevelled with Eph tyrosine kinase receptor and ephrin mediates cell repulsion. EMBO J 22, 847–858.

Taylor, H.B., Khuong, A., Wu, Z., Xu, Q., Morley, R., Gregory, L., Poliakov, A., Taylor, W.R., and Wilkinson, D.G. (2017). Cell segregation and border sharpening by Eph receptor-ephrin-mediated heterotypic repulsion. J R Soc Interface 14, 20170338.

Venkatareddy, M., Cook, L., Abuarquob, K., Verma, R., and Garg, P. (2011). Nephrin regulates lamellipodia formation by assembling a protein complex that includes Ship2, filamin and lamellipodin. PLoS One 6, e28710.

Voltes, A., Hevia, C.F., Engel-Pizcueta, C., Dingare, C., Calzolari, S., Terriente, J., Norden, C., Lecaudey, V., and Pujades, C. (2019). Yap/Taz-TEAD activity links mechanical cues to progenitor cell behavior during zebrafish hindbrain segmentation. Development 146.

Wang, Y., Shang, Y., Li, J., Chen, W., Li, G., Wan, J., Liu, W., and Zhang, M. (2018). Specific Eph receptor-cytoplasmic effector signaling mediated by SAM-SAM domain interactions. Elife 7 e35677.

Wu, Z., Ashlin, T.G., Xu, Q., and Wilkinson, D.G. (2019). Role of forward and reverse signaling in Eph receptor and ephrin mediated cell segregation. Exp Cell Res 381, 57–65.

Yu, J., Peng, H., Ruan, Q., Fatima, A., Getsios, S., and Lavker, R.M. (2010). MicroRNA-205 promotes keratinocyte migration via the lipid phosphatase SHIP2. FASEB J 24, 3950–3959.

Zhuang, G., Hunter, S., Hwang, Y., and Chen, J. (2007). Regulation of EphA2 receptor endocytosis by SHIP2 lipid phosphatase via phosphatidylinositol 3-Kinase-dependent Rac1 activation. J Biol Chem 282, 2683–2694.

Zimmer, M., Palmer, A., Kohler, J., and Klein, R. (2003). EphB-ephrinB bi-directional endocytosis terminates adhesion allowing contact mediated repulsion. Nature Cell Biology 5, 869–878.

